# DNA sequence context controls the binding and processivity of the T7 DNA primase

**DOI:** 10.1101/266924

**Authors:** Ariel Afek, Stefan Ilic, John Horton, David B. Lukatsky, Raluca Gordan, Barak Akabayov

**Author notes:** These authors contributed equally to this work (Ariel Afek, Stefan Ilic).

## Abstract

Primases are key enzymes involved in DNA replication. They act on single-stranded DNA, and catalyze the synthesis of short RNA primers used by DNA polymerases. Here, we investigate the DNA-binding and activity of the bacteriophage T7 primase using a new workflow called <u>H</u>igh-<u>T</u>hroughput <u>P</u>rimase <u>P</u>rofiling (HTPP). Using a unique combination of high-throughput binding assays and biochemical analyses, HTPP reveals a complex landscape of binding specificity and functional activity for the T7 primase, determined by sequences flanking the primase recognition site. We identified specific features, such as G/T-rich flanks, which increase primase-DNA binding up to 10-fold and, surprisingly, also increase the length of newly formed RNA (up to 3-fold). To our knowledge, variability in primer length has not been reported for this primase. We expect that applying HTPP to additional enzymes will reveal new insights into the effects of DNA sequence composition on the DNA recognition and functional activity of primases.

## INTRODUCTION

Replication of chromosomal DNA is one of the central events in every cell cycle. DNA replication is performed by the replisome, a multi-protein complex that includes, among other enzymes, the DNA primase. The primase binds a single stranded DNA (ssDNA) template and synthesizes RNA primers for initiation of the Okazaki fragment formation by the DNA polymerase (Frick and Richardson, 2001). The binding affinity of the primase for its DNA template is based on the recognition of specific DNA sequences along the genome (Soultanas, 2005). For example, the primase domain of gene 4 protein (helicase-primase, gp4) of bacteriophage T7 (referred to simply as the T7 primase), which is the focus of our study, is known to recognize the specific trimer 5’-GTC-3’ (Tabor and Richardson, 1981). After synthesis of tetraribonucleotides (or longer oligos), gp4 must transfer them to the DNA polymerase for use as primers to initiate DNA synthesis (Zhu et al., 2010).

Prokaryotic primases, which include the T7 primase, share high structural similarity and are composed of two highly conserved regions: the N-terminal zinc binding domain (ZBD) (Kato et al., 2003), and the RNA polymerase domain (RPD) (Lee and Richardson, 2005). DNA recognition is mediated by the ZBD, and the condensation of nucleoside triphosphates (NTPs) is catalyzed by the RPD. Currently, it is widely accepted that in bacteriophage T7 the trinucleotide sequence 5’-GTC-3’ is the only preferred site for initiation of primer synthesis (Frick and Richardson, 2001; Lee et al., 2010). However, this trinucleotide sequence is not sufficient to define primase activity. Across the T7 genome, the trinucleotide 5’-GTC-3’ occurs, on average, every 30-bp, with a total of 1,322 occurrences genome-wide. But very few of these sites are actually used by the primase as recognition sites or start sites for RNA primer formation (Frick and Richardson, 2001). The length of the synthesized primers is also known to be important for DNA replication. For example, in the case of the Herpes simplex primase, it was previously demonstrated that although the helicase-primase complex synthesizes primers of 2 to 13 nucleotides, the polymerase only elongates those that are at least 8 nucleotides long (Cavanaugh and Kuchta, 2009). In the case of bacteriophage T7, it is known that although the helicase-primase (gp4) can catalyze the template-directed synthesis of diribonucleotides and triribonucleotides, these short oligonucleotides cannot serve as primers for the T7 DNA polymerase (Kusakabe and Richardson, 1997). Delivery of tetranucleotide primers is sufficient at high concentrations of the RNA primers (Zhu et al., 2010); however, at physiological concentrations, it remains an open question whether longer primers may be required. Swart and Griep (Swart and Griep, 1995) have shown that the DNA primase of E. coli has the ability to produce overlong primers (two to three template lengths) after long incubation times. These overlong primers are not initiated from the recognition sequence (5’-CTG-3’) and are not considered relevant. However, all template length dependent primers are considered biologically significant.

Here, we demonstrate that for bacteriophage T7, DNA sequences flanking the specific recognition trimer 5’-GTC-3’ significantly influence binding preferences and the length of the primers synthesized by the DNA primase. Our finding was possible due to a unique combination of protein-DNA binding experiments and functional activity assays, integrated into a new framework that we call <u>H</u>igh-<u>T</u>hroughput <u>P</u>rimase <u>P</u>rofiling, or HTPP. Our framework can easily be applied to characterize the recognition properties and functional activity of any DNA primase with a known or unknown recognition sequence.

## RESULTS AND DISCUSSION

We developed a new workflow called <u>H</u>igh-<u>T</u>hroughput <u>P</u>rimase <u>P</u>rofiling (HTPP), which combines high-throughput protein-DNA binding measurements and biochemical analyses to define the influence of sequence environment on the DNA-binding and the processivity of DNA primases (**Figure 1**). We performed DNA-binding experiments using custom-designed, single-stranded oligos attached to a glass slide. Similar approaches that had been developed in the past (Berger et al., 2006; Bulyk et al., 2001; Gordan et al., 2013; Warren et al., 2006) were not suitable for characterizing enzymes that bind single-stranded DNA with low affinity, such as DNA primases. Therefore, we have modified both the experimental protocols and the design of the DNA library to suit our purposes (see Experimental Procedures).

We used the T7 DNA primase to develop our assay. The T7 primase serves as an excellent model system because its recognition sequence is known (5’-GTC-3’), and its binding to ssDNA has been previously characterized (Frick and Richardson, 2001). To comprehensively determine the binding specificities of the T7 DNA primase for DNA sequences carrying the recognition site within a wide range of different flanking regions, we designed a custom, single-stranded DNA microarray containing ~29,000 different sequences, with each sequence being present in six replicate spots randomly distributed across the array surface. All sequences are 36-nt long and contain the T7 trinucleotide recognition site 5’-GTC-3’ within flanks with different sequence composition and different symmetries. The recognition sequence was placed at a fixed position at the center of each sequence (**Figure 1A**). The DNA library was commercially synthetized on a microarray slide (Agilent).

The sequence flanks were designed to possess unique sequence features extending beyond the known recognition site. Specifically, we used eight main groups of flanking sequences characterized by a variable nucleotide content: T and G (group 1), T and C (group 2), C and A (group 3), A and G (group 4), G, C and T (group 5), C, T and A (group 6), G, A and T (group 7), and G, A and C (group 8). We designed 2,000-4,000 different sequences for each group. Within each sequence group, we generated subgroups characterized by different symmetries and different numbers of repetitive sequence elements (see examples in **Figure 1A**). The design procedure is conceptually similar to the one described in our past work (Afek et al., 2014). We also included a control sequence group to determine the effect of a single nucleotide adjacent to a specific recognition sequence, 5’-<u>N</u>-GTC-<u>N</u>-3’. In this group, we had 800 sequences with all 16 possible combinations for the two adjacent nucleotides, where each of the 16 variants was embedded in 50 random sequence environments. In order to evaluate the robust ness of the obtained primase-DNA binding preferences and to reduce experimental noise, each sequence was printed in six separate replicate spots on the slide.

**Figure 1.**
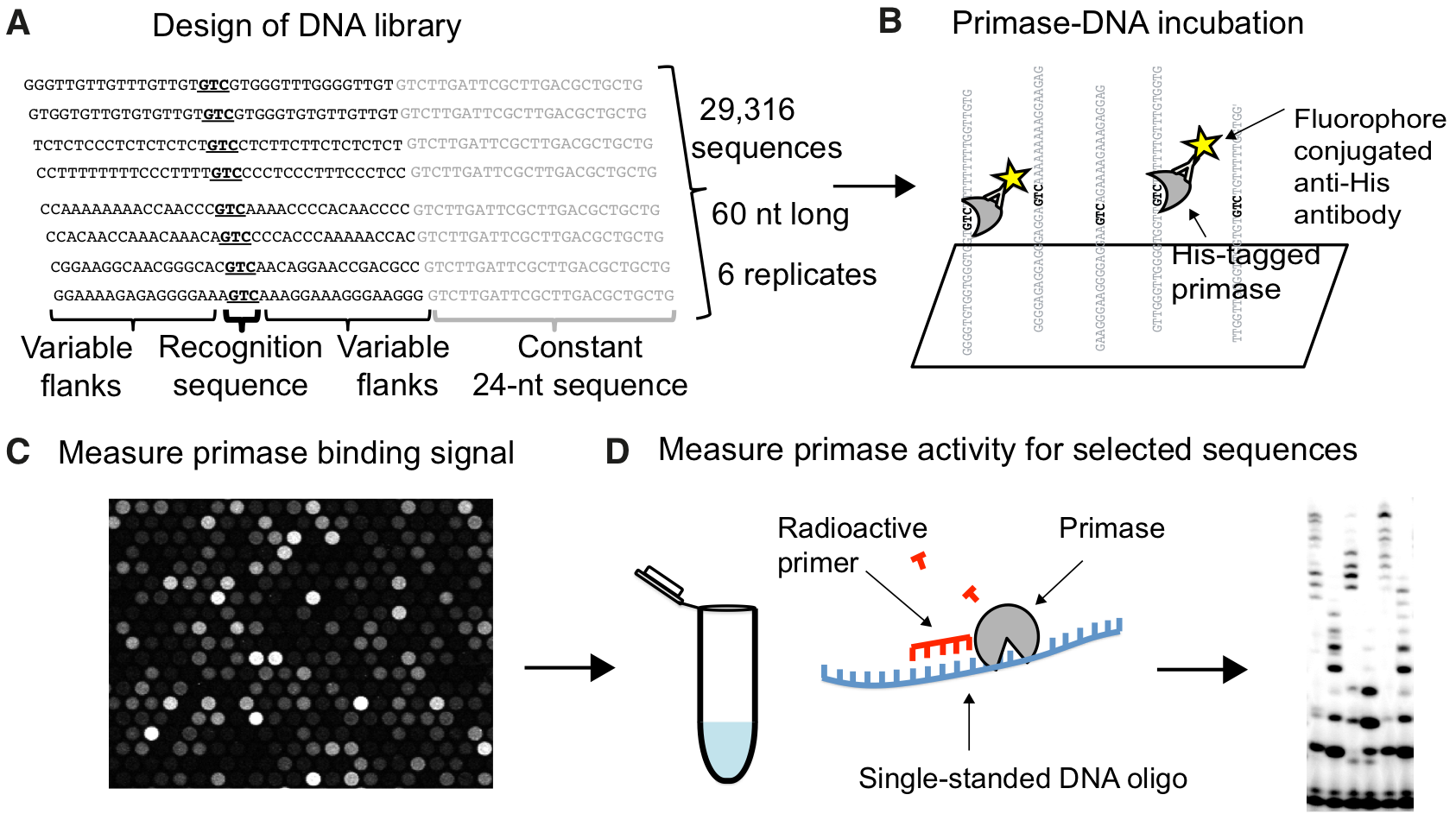
High-Throughput Primase Profiling (HTPP) workflow. **(A)** Design of DNA sequence library for the T7 primase: 29,316 unique sequences; 6 replicates/sequence. Probes contain custom 36-nucleotide sequences containing the recognition site (GTC) within variable flanking regions, followed by a constant 24-nucleotide sequence tethered to a glass slide. **(B)** Schematic representation of primase incubation and detection on the slide. **(C)** The slide was scanned to detect protein binding, as indicated by Alexa 488-labeled anti-His antibodies. Image shows magnification of a portion of the scan. The strength of primase binding to the single-stranded DNA molecules in each spot is represented by the fluorescence intensity of the spot, ranging from highest/saturated intensity (white) to no signal (black). **(D)** DNA sequences with very high or very low primase binding intensity were selected and biochemically tested for activity by primer synthesis assay.

The measured primase-DNA binding preferences exhibit a wide range of intensities for the different groups of sequences (**Figure 2**). The nucleotide composition of the flanking DNA regions was a major factor determining the fluorescence signal caused by primase-DNA binding. We emphasize that the observed variability in the measured fluorescence intensity signal cannot be explained solely based on the effect of nucleotides adjacent to the motif (5’-<u>N</u>-GTC-<u>N</u>-3’), as demonstrated by the control group of sequences (**Figure S1A-D**). Thus, we conclude that the composition of flanking DNA regions outside of the specific recognition site significantly affects primase-DNA binding preferences in a highly non-random manner (**Figure 2A**). For example, changing the entire flanking composition from T/G to A/G increases the measured florescence intensity by ~5 fold (on average). As shown in **Figure 2B**, the binding distributions are significantly different between sequences with T/G versus A/G composition (p-value < 10e-323). For a randomly selected pair of sequences, the binding affinity difference between the T/G and the A/G contexts was validated by gel retardation assay (**Figure S2**).

In addition to the sequence composition of the flanking regions, their symmetry is also an important determinant of DNA binding. Within each group of probes with a specific sequence composition, our design includes sequences with different arrangements of the nucleotides. For example, GGGTTT and GTGTGT have the same composition but different symmetries.

**Figure 2.**
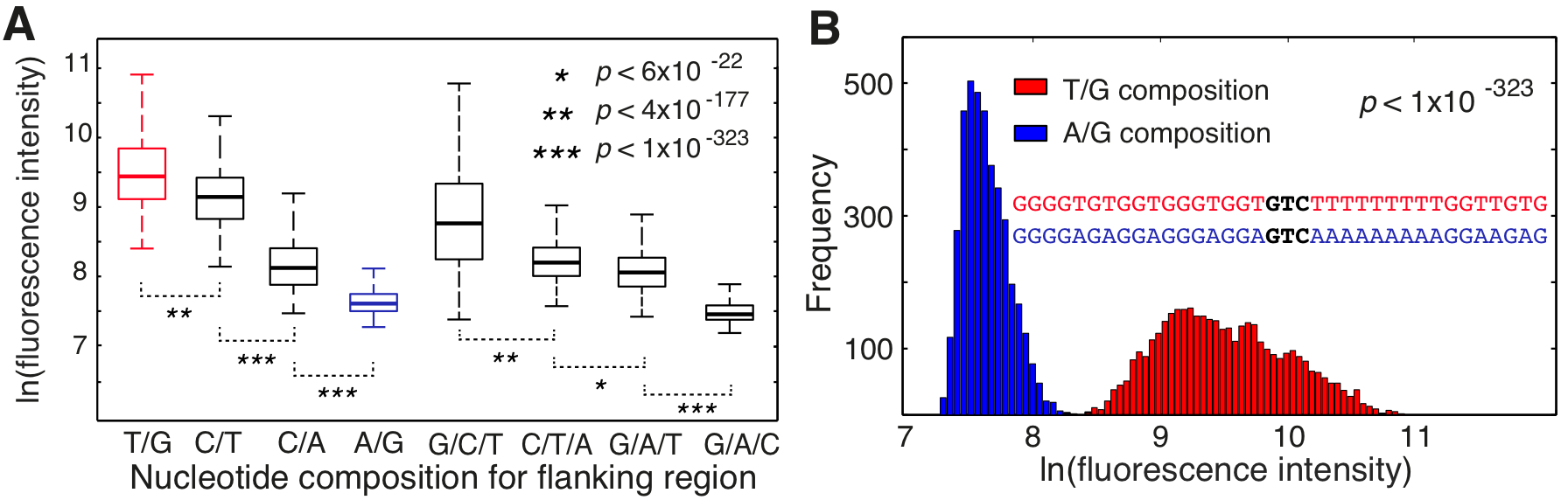
Effect of sequence context on the DNA-binding specificity of the T7 primase. **(A)** Distribution of measured binding intensities for DNA probes containing the recognition trinucleotide 5’-GTC-3’ flanked by sequences with different nucleotide composition (one-way ANOVA p-values are shown). **(B)** Primase binding intensities for oligonucleotides with T/G versus A/G sequence compositions are highly significantly different (p-value < 10^−323^; one-way ANOVA). Inset shows an example from each sequence group.

We have previously demonstrated that these symmetries have a major influence on transcription factor-DNA binding (Afek et al., 2014). Thus, we asked whether DNA repeat symmetries also influence the DNA binding preferences of primases. Indeed, for the T7 primase tested in our study we observed a strong correlation between the number of repetitive DNA sequence elements and the primase binding affinity (**Figure S1E-G**). Taken together, our results demonstrate a complex landscape of primase binding preferences to DNA that extends far beyond a classical picture of the specific 5’-<u>N</u>-GTC-<u>N</u>-3’ recognition sequence. This landscape is affected by both the DNA composition and the DNA repeat symmetry.

Next, we investigated whether the measured primase-DNA binding preferences correlate with the functional, biochemical activity of the T7 primase. We selected 20 representative DNA sequence pairs (i.e. DNA templates) (**Figure S3A**), from two groups with significantly different primase binding signals (**Figure 2B**, **Figure S3B**). The first group contained DNA sequences with T/G enriched flanks, and the second group contained DNA sequences with A/G enriched flanks. These two groups of sequences exhibit very different binding profiles, with significantly stronger primase binding to the T/G group as compared with the A/G group (**Figure 2B**, **Figure S3B**). Next, we compared the RNA primer products for each DNA template from the T/G group to its matching sequence from the A/G group, using primer synthesis assays.

Interestingly, longer RNA primers were formed from the DNA templates characterized by higher affinity of primase-DNA binding (**Figure 3A,B**). Our results for the 20 selected pairs of templates show that changing the flanking regions from A/G-rich to T/G-rich generally results in the production of longer primers, with an increase in the maximum primer length of as much as 200% (e.g. from 7 to 21 nucleotides, for sequence pair 17) (**Figure 3A**). These results indicate a higher processivity of the DNA primase for the higher affinity, i.e. for the T/G-rich, DNA templates. Processivity is usually defined as the number of incorporations of nucleoside monophosphate to the growing primer until the enzyme dissociates from its DNA track. So far, the processivity has been considered an internal property of the enzyme, and the effect of DNA sequence on the processivity of polymerases is usually neglected (Von Hippel et al., 1994). In particular, highly processive enzymes are characterized by large and hydrophobic DNA-binding surfaces (Akabayov et al., 2010). A larger DNA-binding surface of the DNA polymerase has been shown to increase its affinity for DNA and its ability to better slide on the DNA (Akabayov et al., 2010). Our current results show, for the first time, that DNA sequence also has a significant effect on primase processivity. Specifically, our results suggest that the processivity of the T7 primase depends on the affinity of the primase for the DNA template, with DNA sequences characterized by stronger primase binding promoting the production of longer RNA primers (**Figure 3**, **Figure S3**).

**Figure 3.**
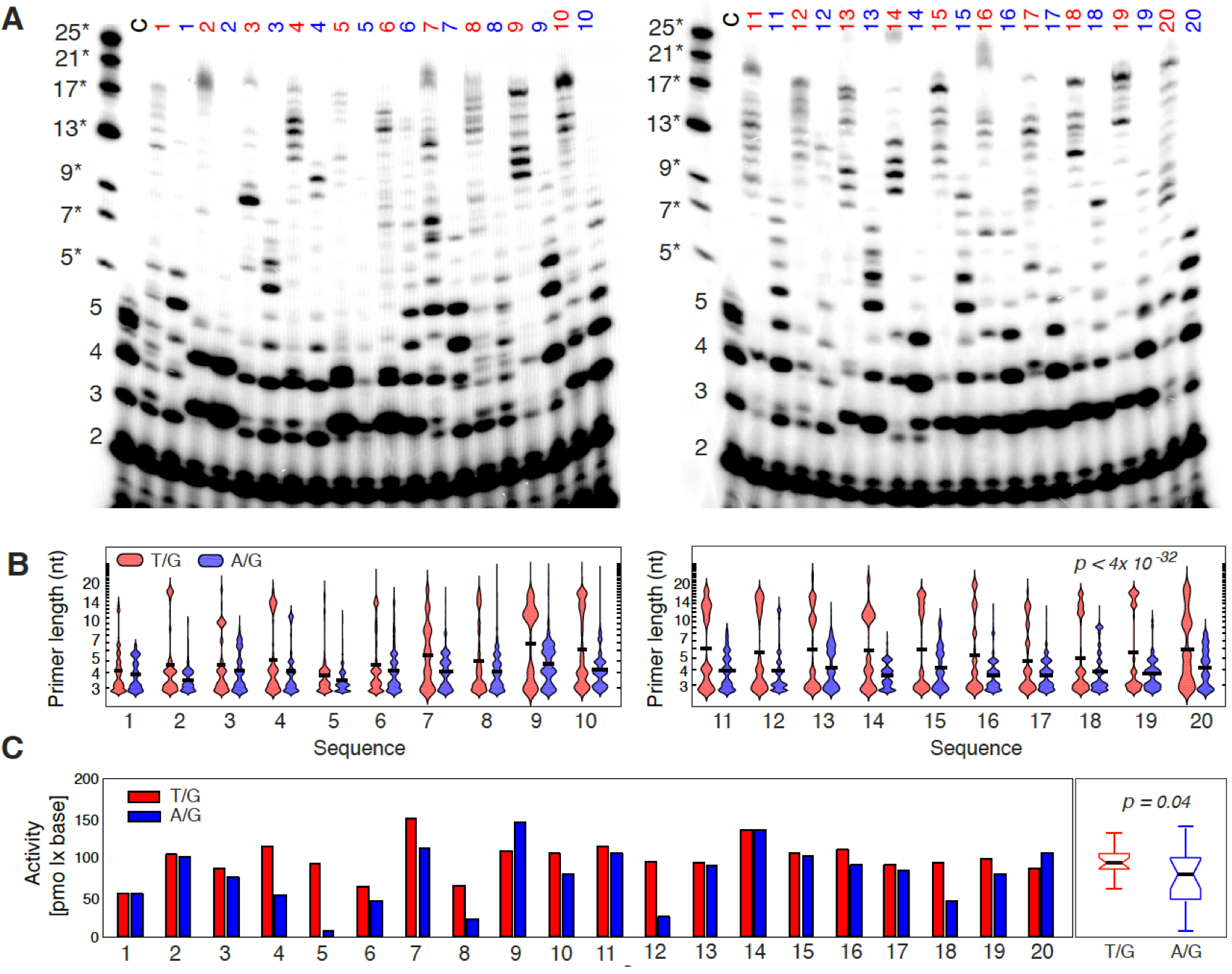
Template-directed RNA primer synthesis catalyzed by the T7 DNA primase. **(A)** Oligonucleotide synthesis by the T7 DNA primase. The reactions contained oligonucleotides with the primase recognition sequence (numbered lanes), a positive control oligonucleotide 5’-GGGTCA_10_-3’ (lane “C”), and ^32^P-γ-ATP, ATP, CTP, UTP, and GTP in the standard reaction mixture. After incubation, the radioactive products were analyzed by electrophoresis through a 25% polyacrylamide gel containing 7 M urea, and visualized using autoradiography. Size marker (left-most lane) contained 7 short commercially synthesized RNA molecules (size range 5-25 nt), which we denote 5* to 25*. These molecules have a single phosphate group at the 5’ end, and are thus different from the RNA primers synthesized by T7 primase, which contain three negatively charged phosphate groups at the 5’ end, originating from the first incoming ATP. For this reason, as shown in the gel image, the retardation of the 5-mer RNA primer synthesized by T7 primase (lane C) was faster compared to the commercially synthesized 5-mer RNA primer with only one phosphate group at its 5’ end. **(B)** RNA primer length distributions. Violin plots show the probability densities of primer fragment length for the 20 T/G and A/G sequence pairs, as quantified from the gel using Matlab. Primer length was estimated from the size markers and the control RNA primers formed on 5’-GGGTCA_10_-3’ template. All primers of three or more nucleotides are shown. The violin plots demonstrate that, for all 20 pairs tested, longer RNA primers are formed for the templates with T/G versus A/G contexts (Wilcoxon rank sum test p-value < 3e-28). **(C)** Quantification of T7 primase activity. Left plot shows the normalized activity for 20 pairs of DNA sequences, where one sequence contains Ts (red) and the second sequence has all the Ts replaced by As (except for the T in the recognition sequence GTC). The normalized activity was computed as a sum of the products between the total intensity of the gel bands (in picomols quantified by autoradiography) and the primer lengths (estimated from the size markers and the control RNA primers). Right boxplot compares the primase activity for sequences containing T/G (red) or A/G (blue) in the flanking regions of the T7 DNA primase recognition sequence, combined over all 20 pairs. The difference between the two groups is statistically significant (p-value = 0.04, one-way ANOVA). All primers of four nucleotides or more were included in this analysis, as it has been previously shown that primers as short as four nucleotides can be functional (Zhu et al., 2010). Including di- and trinucleotides changes the trend; however, such short oligoribonucleotides can-not serve as functional primers for the T7 DNA polymerase (Kusakabe and Richardson, 1997).

In order to characterize the effect of flanking sequences on primase activity, we quantified and compared the activity, expressed as pmol x base, for the 20 selected template pairs (**Figure 3C**) taking into account RNA oligos of four nucleotides or longer, which can serve as functional primers for T7 DNA polymerase (Romano and Richardson, 1979). For the T7 primase, it is known that ATP is always the first incoming nucleotide. We used [γ–^32^P] ATP to 5’ end-label the RNA primers, which ensured that each primer was labeled only once and thus allowed us to quantify the absolute amount of RNA primers (**Figure 3**). For all 20 templates, we found that replacing the T’s in the flanks with A’s reduces the processivity of the primase (**Figure 3A**, **Figure S4A**), consistent with the lower binding affinity measured in our high-throughput assay (**Figure S3B**). These results suggest that, in general, higher binding affinity for the DNA recognition sequence yields a higher processivity of the T7 DNA primase, a feature that has not been explored thus far. We have validated these finding by repeating the experiment with the full-length gp4 protein (**Figure S3C**). Interestingly, the total amount of RNA molecules formed (across fragments of all lengths) is not higher for the high affinity templates (**Figure S3D**). However, the processivity differs significantly, with high affinity templates leading to formation of longer primers (**Figures S3C**, **S4B**).

In summary, it has long been assumed that primase start sites are dictated by the presence of 5’-GTC-3’ trinucleotides. Here, we show that this trinucleotide sequence is not sufficient to define the recognition of DNA templates by the T7 primase. In particular, we show that both the sequence composition and the DNA repeat symmetry in DNA regions flanking the specific recognition site significantly influence primase-DNA binding preferences. We also demonstrate that these binding preferences are correlated with the functional enzymatic activity. The fact that recognition sequences with higher binding affinity lead to formation of longer primers is significant. At the high-affinity sites, the preference for forming longer primers will lead to a decreased probability of forming short, futile primers from the DNA template. We note that primers as short as four nucleotides can be functional; Richardson and co-workers have shown that delivery of tetranucleotide primers from the primase to the DNA polymerase is possible at high concentrations of the RNA primers (Zhu et al., 2010). However, at physiological concentrations, longer primers may be more important for effective delivery to the DNA polymerase.

Our array-based assay provides, for the first time, a fast, cheap, and high-throughput way to determine the DNA recognition properties of primases. The assay can be used with any recombinant primase protein, and can accommodate almost 30,000 DNA template sequences of up to 60 nucleotides each. We anticipate that applying this assay to more primase enzymes and DNA templates will reveal complex DNA recognition patterns. Our binding assay does not directly assess the processivity/activity of the tested enzyme. However, due to its high throughput and highly quantitative nature, the array-based binding assay provides an unbiased way to test a very large number of sequences in order to identify the best candidates for further biochemical characterization. We anticipate that the combination of these two complementary approaches, which we refer to as the HTPP framework, will lead to new insights into the DNA sequence features important for the functional activity of primases, such as extended primase recognition patterns and influences from sequence content and sequence symmetry on the binding properties and the processivity of DNA primases.

## EXPERIMENTAL PROCEDURES

### Design of the DNA library

We designed a custom DNA library containing 29,316 distinct sequences centered at the putative recognition site for the T7 DNA primase (5’-GTC-3’). Each sequence is composed of 60 nucleotides and has the general form (N)_17_ GTC (N)_16_-GTCTTGATTCGCTTGACGCTGCTG. The recognition site (GTC) and the last 24 nucleotides (closest to the glass slide) were kept constant. (N)_17_ and (N)_16_ corresponds to the variable sequences flanking the recognition site in each probe.

The flanks were designed to have unique context features extending beyond the known recognition site. One such feature is the nucleotide composition, which we chose to represent in eight main categories of different flanking sequences composed of 2 or 3 specific nucleotides, as follows: T and G (group 1); T and C (group 2); C and A (group 3); A and G (group 4); G, C and T (group 5); C, T and A (group 6); G, A and T (group 7); G, A and C (group 8). We designed 2,000 different sequences for each group. Within each composition group, we had subgroups of sequences with different types and different numbers of sequence repeats.

Finally, we added a control group to determine the effect of the nearest nucleotide on primase binding to the conserved recognition sequence. The control group contained 800 sequences with all 16 possible combinations for the trimer neighbors 5’-N-GTC-N-3’, with each combination present in 50 random environments.

Although our binding assay was adapted after the protein-binding microarray (PBM) technology of (Berger et al., 2006), the DNA library design used in this study was very different from the widely used universal PBM design (Berger and Bulyk, 2009; Berger et al., 2006). The universal PBM design can be used to determine the average binding specificity of a protein for all possible 8-bp sequences. In contrast, our custom library was based on DNA sequences with a central GTC core flanked by unique sequence elements specifically designed to enable us to detect long-range effects, which are impossible to detect using a universal PBM library.

### HTPP (High-Throughput Primase Profiling) assay

The custom DNA library containing 29,316 template sequences was synthetized *de novo*, as single-stranded DNA molecules, on a microarray slide (Agilent Technologies). We used the 4×180K Agilent microarray format (AMADID #78366), which allowed each sequence of interest to be represented in 6 replicate DNA spots, randomly distributed across the slide. We pre-wet the single-stranded DNA microarray in PBS and 0.01% Triton X-100, then blocked it with 2% milk in PBS for 1 hour. We washed the blocked microarray once with PBS/ 0.1% Tween 20 and once with PBS/0.01% Triton X-100, similar to the protein-DNA binding microarray protocol of (Berger and Bulyk, 2009). After the blocking step, the microarray was incubated, for 30 minutes, with a PBS-based protein binding mixture of 5 μM His-tagged protein, 6.5mM MgCl_2_, 30mM KGlu, 6mM DTT, 65μM rNTP (all four), 2% milk, 100 ng/μL BSA, 50 ng/μL Salmon Testes DNA, and 0.02% TX-100. After incubation, the microarray slide was briefly rinsed in PBS. Next, the bound protein was tagged with 10 ng/μL anti-His antibody conjugated to Alexa 488 (Qiagen; 35310) in PBS with 2% milk and binding buffer, for 30 minutes. Finally, the microarray slide was briefly rinsed with PBS. Compared to the universal PBM protocol of (Berger and Bulyk, 2009), we have reduced the duration of the protein and antibody incubation steps, we substantially changed the binding and antibody buffer conditions, and we reduced or eliminated the harsh washing steps, since they were not suitable for low affinity enzymes such as the T7 primase. Microarray scanning and quantification were performed using a GenePix 4400A scanner (Molecular Devices), and the data were analyzed using custom scripts in order to obtain fluorescence intensities for all sequences represented on the array. Each sequence was present in six replicate spots in order to measure the robustness of the results and to reduce experimental noise. For each sequence, we report the median fluorescence intensity over the six replicates.

### Gel retardation assay

T7 DNA primase in storage buffer (50mM Tris-HCl, pH 7.5; 1mM DTT, 1mM EDTA, 50% glycerol) was equilibrated with DNA for 20 minutes on ice. The buffer composition of the gel retardation assay was optimized to obtain the maximum resolution for resolving DNA. Reactions (final volume 10 μL) were resolved by electrophoresis at 4°C through native gel containing 6% polyacrylamide (29:1 acrylamide: bisacrylamide) in 1 × TBE buffer. DNA concentration was 15 nmol, and T7 primase was added to a final concentration of 0, 5.4, 16, 49, 145, 440 μM, respectively. Autoradiographs of the dried gels were analyzed by densitometry using Fujifilm PhosphorImager.

### Materials

Chemicals were purchased from Sigma. ATP and CTP were purchased from Roche Molecular Biochemicals. [α-^33^P] CTP and [ γ-^32^P] ATP (3000 Ci/mmol) were purchased from Perkin Elmer.

### Protein expression and purification

T7 primase domain (residue: 1-271) was over-produced and purified as previously described (Zhu et al., 2009). Recombinant gene 4 protein (gp4) was over-produced and purified as previously described (Lee and Richardson, 2001). All chemical reagents were of molecular biology grade (Sigma); ATP and CTP (Roche Molecular Biochemicals).

### Oligoribonucleotide synthesis assay by T7 DNA primase

Oligoribonucleotides were synthesized by DNA primase. And the product was measured as previously described (Ilic et al., 2016), in reactions containing 1 μM of gene 4 primase domain. Standard 10 μL reaction contained 5 μM of DNA template (5’-GGGTCA_10_-3’ was the DNA recognition sequence in the control reaction), 200 μM rNTPs, 200 μM [α-^32^P]-CTP, and primase domain in a buffer containing 40 mM Tris-HCl (pH 7.5), 10 mM MnCl_2_, 10 mM DTT, and 50 mM KGlu. After incubation at room temperature for 20 minutes, the reaction was terminated by adding an equal volume of sequencing buffer containing 98% formamide, 0.1% bromophenolblue, and 20 mM EDTA. The samples were loaded onto a 25% polyacrylamide sequencing gel containing 7 M urea, and visualized using autoradiography.

### Oligoribonucleotide synthesis assay by full-length gp4

Oligoribonucleotides were synthesized in reactions containing 500 nM of gene 4 protein. Standard 10 μL reaction contained 20 μM of DNA template, 500 μM rNTPs, 500 μM [α–^32^P]-ATP, and gp4 protein in a buffer containing 40 mM Tris-HCl (pH 7.5), 10 mM MnCl_2_, 10 mM DTT, and 50 mM KGlu. After incubation at 37°C for 30 minutes, the reaction was terminated by adding an equal volume of sequencing buffer (98% formamide, 0.1% bromophenolblue, 20 mM EDTA). The samples were loaded onto a 25% polyacrylamide sequencing gel containing 7 M urea, and visualized using autoradiography.

## SUPPORTING INFORMATION

**Figure S1.**
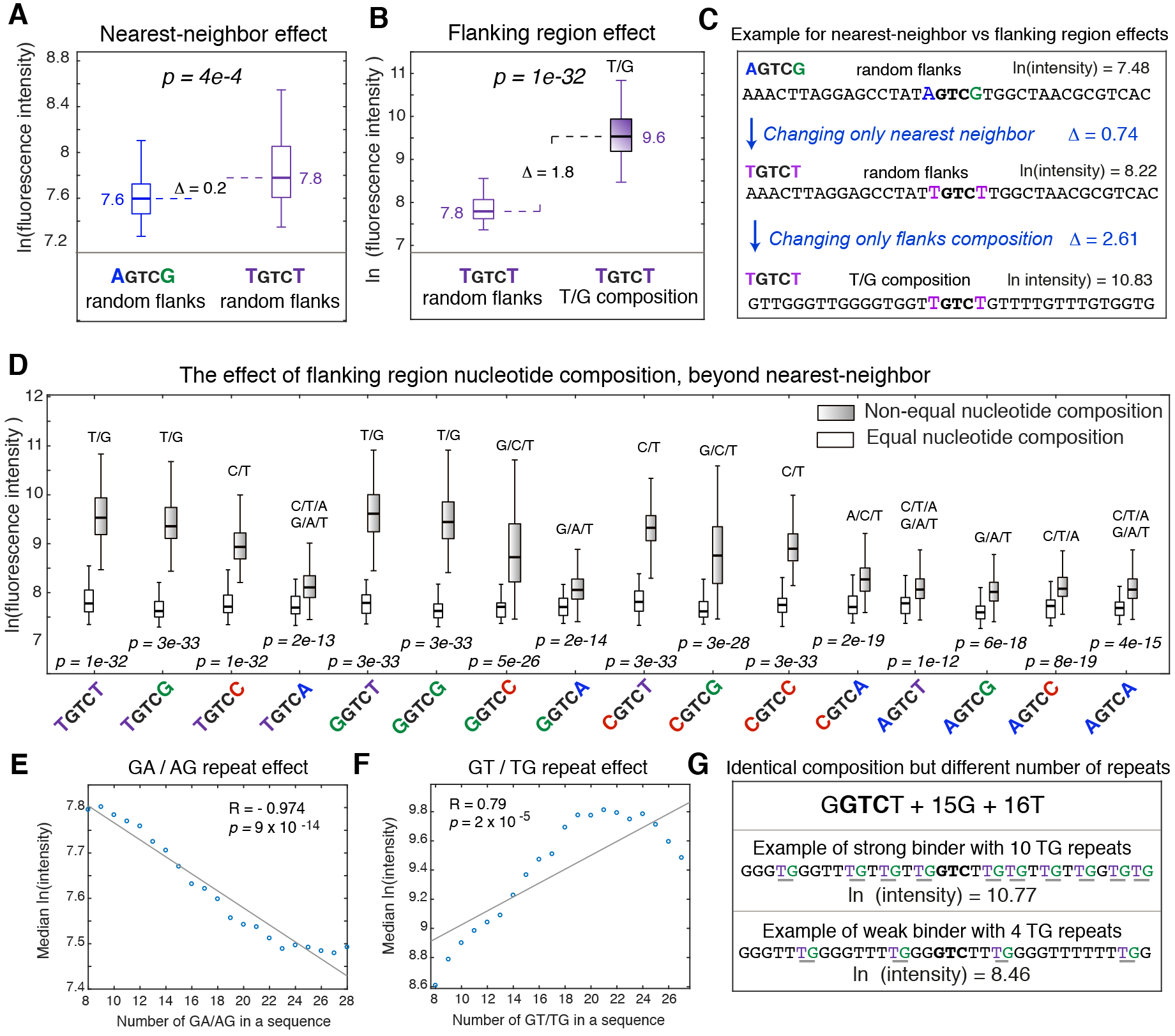
The effects of different sequence features on T7 primase binding to DNA. **(A)** Boxplot showing the median fluorescence intensity and the distribution of intensities for two sets of 50 identical random sequences, which differ only by nearest nucleotides flanking the 5’-GTC-3’ core. These two sets represent the largest and most significant difference detected for nearest-neighbor nucleotide effects (Wilcoxon rank sum test p-value = 4e-4). **(B)** Boxplot showing the median fluorescence intensity and the distribution of intensities for sequences containing T as the nearest-neighbor nucleotide, and either G/T or random sequence in the rest of the flanking regions. Here, the binding changes significantly due to distal flanking region composition (Wilcoxon rank sum test p-value = 1e-32). The effect of sequence composition (shown in panel B) is considerably stronger compared to nearest-neighbor effect (shown in panel A). **(C)** Two examples of binding signal, for sequences from the two groups shown in panels A and B. **(D)** Comparison of fluorescence intensity distribution, for all 16 possibilities of nucleotides adjacent to the 5’-GTC-3’ core, in two different compositional environments (random vs. non-random). For all cases we see that the flanking sequence composition can significantly increase primase binding (Wilcoxon rank sum test p-value < 1e-12). **(E)** The effect of sequence symmetry on primase-DNA binding: higher number of GA/AG repeats within flanking sequences having the same composition (G/A) reduces primase binding to the DNA. **(F)** Higher number of GT/TG repeats within flanking sequences with the same composition (G/T) significantly influences primase binding to the DNA. Interestingly, the measured fluorescence intensity exhibits a non-linear behavior as a function of the number of GT/TG repeats. The intensity peaks at ~20-24 repeats, and decrease beyond that point. **(G)** Two examples of binding signal, for sequences from the same composition group (G/T composition) with the same nucleotides adjacent to the GTC core (both G**GTC**T), but with different numbers of TG dinucleotides in the flanking sequence. The sequence with the larger number of TG repeats has ~10 fold higher fluorescence intensity in the HTPP measurements.

**Figure S2.**
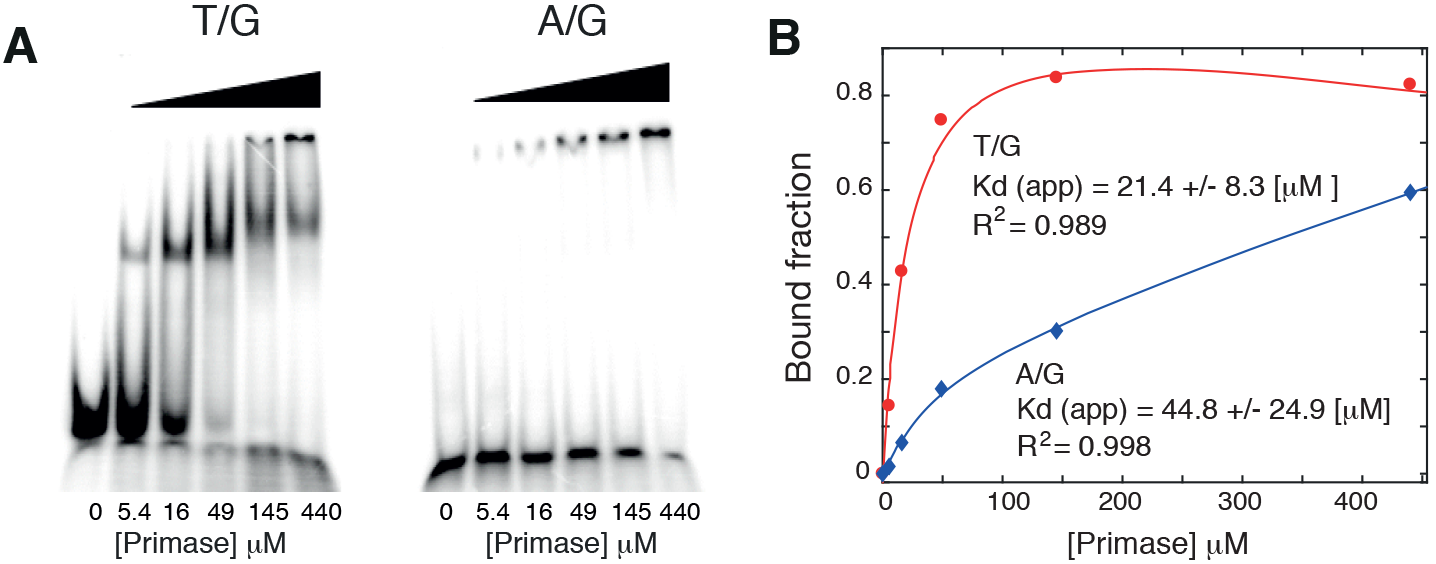
Biochemical analysis for representative DNA sequences. **(A)** Gel retardation assay of selected ^32^P-5’-end-labeled DNA template as a function of increasing concentration of T7 DNA primase, for sequence #11 (from Figure S3) with T/G or A/G composition. **(B)** Binding curves for the T7 primase based on the gel from panel A. Red: T/G rich sequence. Blue: A/G rich sequence. Apparent Kd was estimated using GraphPad Prism 7 (total and nonspecific, one site binding fit, y= Bx/(Kd+x)+Nx+D).

**Figure S3.**
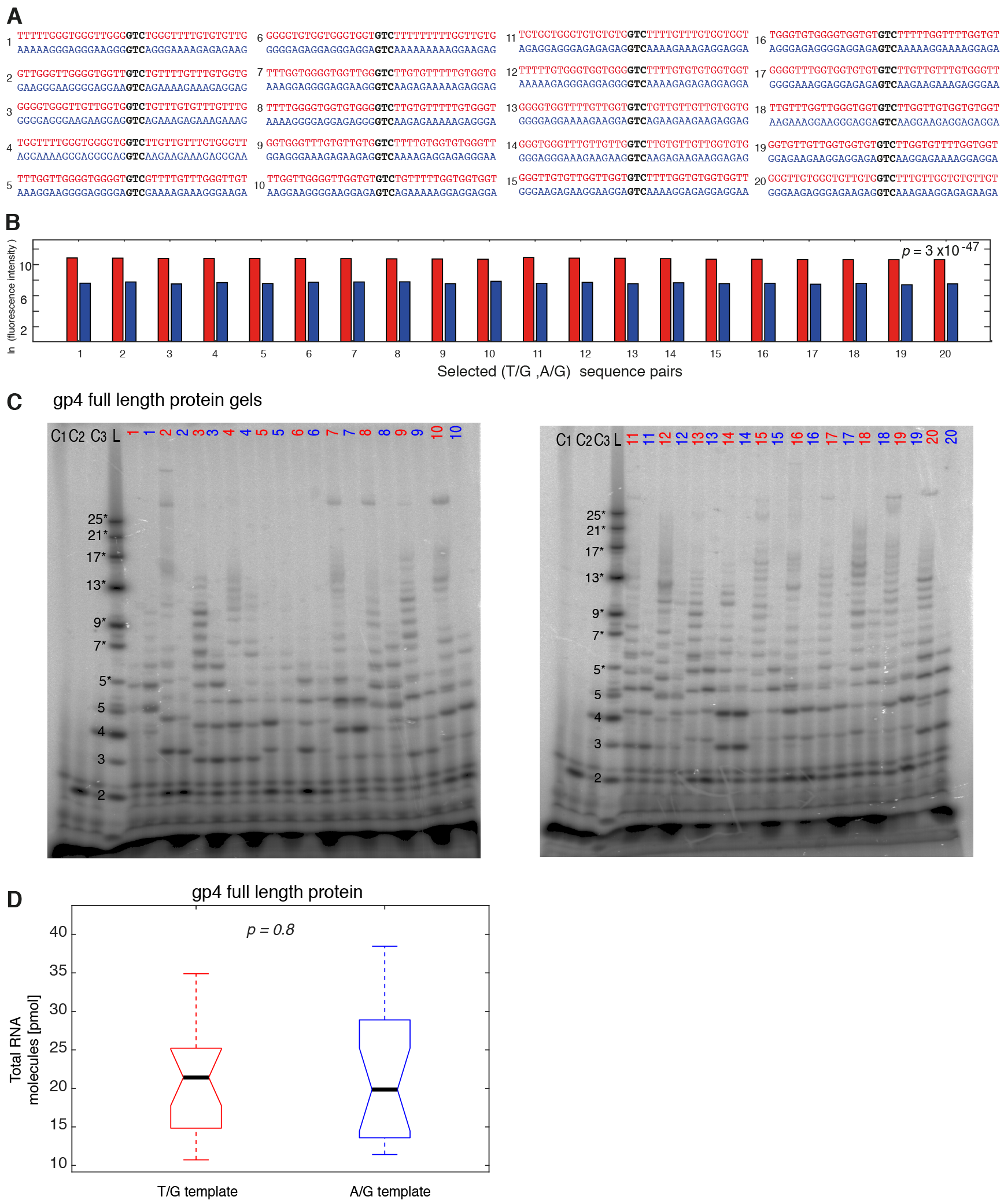
Biochemical analysis for selected DNA sequences. **(A)** The 20 sequence pairs selected for RNA primer synthesis catalyzed by the T7 DNA primase, from two different compositional groups (T/G and A/G). **(B)** Fluorescence intensity measured for the 20 oligo pairs from panel A. **(C)** Template-directed RNA primers synthesis catalyzed by the gp4 full-length protein. The reaction contained oligonucleotides with the primase recognition sequence, two control oligonucleotides: 5’-GGGTCAA-3’ (C2) and 5’-GGGTCA_10_-3’ (C3), and [α-^32^P]-ATP, CTP, GTP, and UTP in the standard reaction mixture. After incubation, the radioactive products were analyzed by electrophoresis through a 25% polyacrylamide gel containing 7 M urea, and visualized using autoradiography. RNA ladder (marked L) contained 7 short commercially synthesized RNA molecules (size range 5-25 nt), which we denote 5* to 25*. These molecules have a single phosphate group at the 5’ end, and are thus different from the RNA primers synthesized by T7 primase, which contain three negatively charged phosphate groups at the 5’ end, originating from the first incoming ATP. For this reason, the retardation of the 5-mer RNA primer synthesized by T7 primase is faster compared to the commercially synthesized 5-mer RNA primer with only one phosphate group at its 5’ end. C1 contains reaction mixture as a negative control. **(D)** Boxplot of the total RNA primer initiation for sequences containing T/G (red) or A/G (blue) in the flanking regions of the gp4 full length protein, combined over all 20 pairs. The difference between the two groups is not statistically significant (p-value = 0.8, one-way ANOVA).

**Figure S4.**
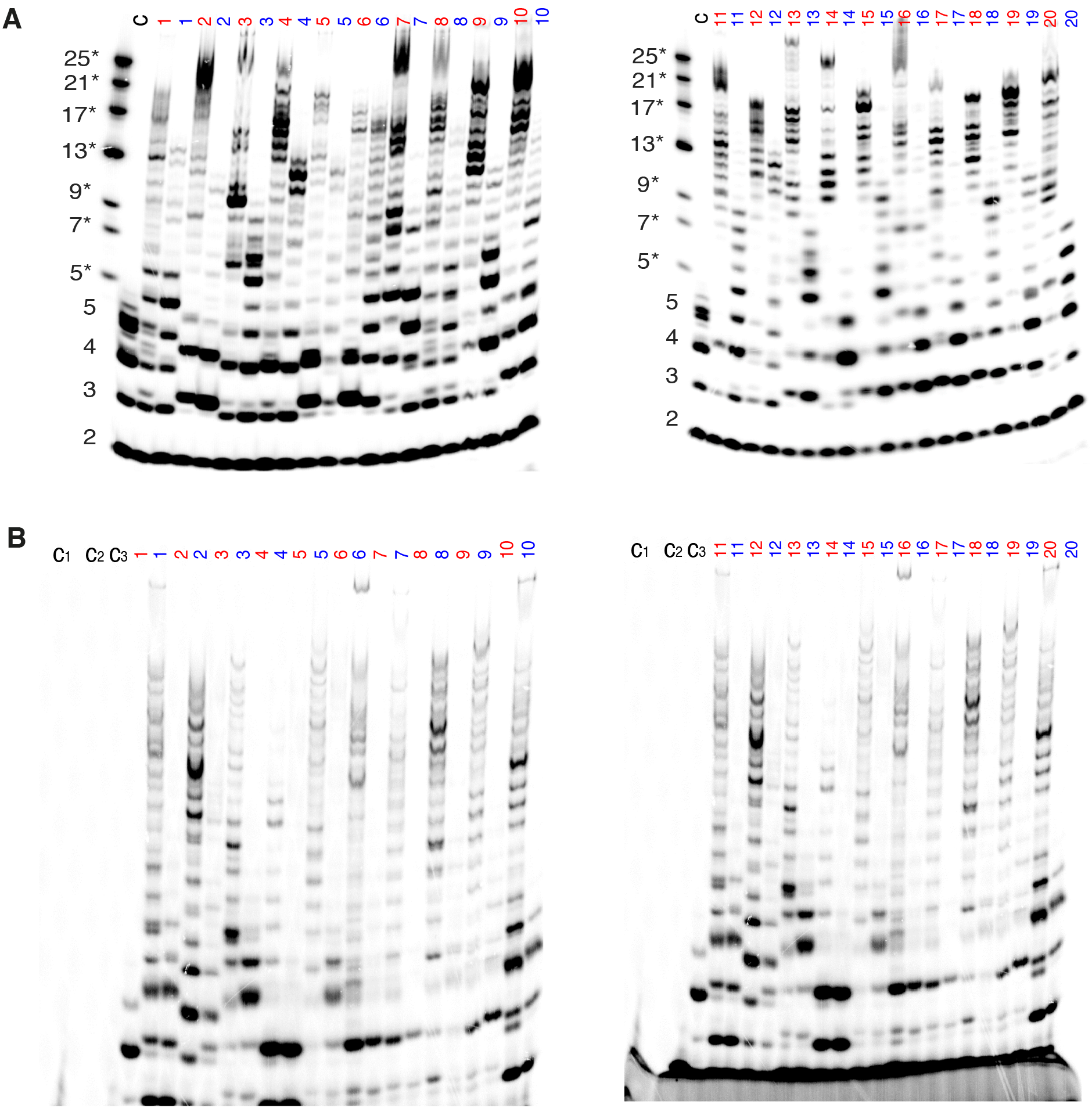
Template-directed RNA primers synthesis. **(A) Catalyzed by T7 DNA primase.** The reaction contained oligonucleotides with the primase recognition sequence, a control oligonucleotide 5’-GGGTCA_10_-3’, and α-^33^P-CTP, ATP, GTP, and UTP in the standard reaction mixture. After incubation, the radioactive products were analyzed by electrophoresis through a 25% polyacrylamide gel containing 7 M urea, and visualized using autoradiography. Size marker (left-most lane) contained 7 short commercially synthesized RNA molecules (size range 5-25 nt), which we denote 5* to 25*. These molecules have a single phosphate group at the 5’ end, and are thus different from the RNA primers synthesized by T7 primase, which contain three negatively charged phosphate groups at the 5’ end, originating from the first incoming ATP. For this reason, as shown in the gel image, the retardation of the 5-mer RNA primer synthesized by T7 primase (lane C) was faster compared to the commercially synthesized 5-mer RNA primer with only one phosphate group at its 5’ end. This figure is similar to Figure 3A, however the signal comes 33P-CMP incorporated into the elongated primer, and not from 32P-ATP end label of the first incoming nucleotide (which is the case in Figure 3A). **(B) Catalyzed by the gp4 full-length protein.** The reaction contained oligonucleotides with the primase recognition sequence, two control oligonucleotides: 5’-GGGTCAA-3’ (C2) and 5’-GGGTCA_10_-3’ (C3), and [α-^33^P]-CTP, ATP, GTP, and UTP in the standard reaction mixture. C1 serves a negative control (only reaction mixture). After incubation, the radioactive products were analyzed by electrophoresis through a 25% polyacrylamide gel containing 7 M urea, and visualized using autoradiography.

